# Common methods for fecal sample storage in field studies yield consistent signatures of individual identity in microbiome sequencing data

**DOI:** 10.1101/038844

**Authors:** Ran Blekhman, Karen Tang, Elizabeth A. Archie, Luis B. Barreiro, Zachary P. Johnson, Mark E. Wilson, Jordan Kohn, Michael L. Yuan, Laurence Gesquiere, Laura E. Grieneisen, Jenny Tung

## Abstract

Field studies of wild vertebrates are frequently associated with extensive collections of banked fecal samples, which are often collected from known individuals and sometimes also sampled longitudinally across time. Such collections represent unique resources for understanding ecological, behavioral, and phylogenetic effects on the gut microbiome, especially for species of particular conservation concern. However, we do not understand whether sample storage methods confound the ability to investigate interindividual variation in gut microbiome profiles. This uncertainty arises in part because comparisons across storage methods to date generally include only a few (≤5) individuals, or analyze pooled samples. Here, we used n=52 samples from 13 rhesus macaque individuals to compare immediate freezing, the gold standard of preservation, to three methods commonly used in vertebrate field studies: storage in ethanol, lyophilization following ethanol storage, and storage in RNAlater. We found that the signature of individual identity consistently outweighed storage effects: alpha diversity and beta diversity measures were significantly correlated across methods, and while samples often clustered by donor, they never clustered by storage method. Provided that all analyzed samples are stored the same way, banked fecal samples therefore appear highly suitable for investigating variation in gut microbiota. Our results open the door to a much-expanded perspective on variation in the gut microbiome across species and ecological contexts.

## Importance

Although variation in gut microbiome profiles is extensively studied, we know little about how this variation is influenced by sample storage and handling. This is especially important for sample collections from field studies, which can be hugely informative resources for microbiome studies, but often utilize variable storage approaches. Here, we compare four fecal sample storage methods that are commonly used in field studies, including freezing, lyophilization, storage in ethanol, and RNAlater. We find that the effect of storage method on microbiome profiles is consistently smaller than the effect of individual identity. Our results indicate that sample storage method is unlikely to affect the results of a study, as long as the same storage method is used for all samples. By indicating the utility of using previously collected sample banks for gut microbiome profiling, our results open the door to a vastly expanded perspective on gut microbiome variation in the natural world.

## Main text

Noninvasive collection is often the only feasible approach for obtaining samples from wild vertebrates, especially in threatened or endangered species [1]. Fecal samples are especially common, as they can be collected without disrupting study subjects, can often be unambiguously assigned to donors, and can be longitudinally collected from the same animal over time. Such samples also contain abundant information about the genetics, endocrinology, and parasite burden of the animals from which they are obtained. For these reasons, fecal samples may be the most extensively banked sample type available for wild vertebrates.

Such collections represent potentially invaluable resources for understanding interindividual variation in the gut microbiome in comparative or conservation contexts. However, sample storage methods vary widely across studies, and in most cases, samples were not collected with microbiome analyses in mind. To assess the potential for mining existing sample banks, we investigated how three common field storage methods affect gut microbiome diversity and composition estimates, compared to the gold standard of immediate freezing. We were particularly interested in comparing the roles of storage method versus individual identity. Although storage methods often explain substantial variation in microbiome composition when all other sources of variance are controlled (e.g., [2, 3], the degree to which they confound other analyses depends on their importance *relative* to the effects of biologically interesting variation (interindividual, temporal, and environmental). Previous studies have focused on small numbers of study subjects (n≤5), limiting their ability to evaluate this question [2-9].

Here, we compared fecal samples collected from 13 captive adult rhesus macaques (*Macaca mulatta*). Each fecal sample was divided into four aliquots (n=52 samples; Supplementary Information, Table S1), stored via: 1) immediate freezing at −20 °C; 2) immersion in absolute ethanol; 3) immersion in the preservative RNAlater; or 4) immersion in ethanol followed by lyophilization to powder (often used for steroid hormone analysis: [10]; drying protocols are also sometimes used for genetic samples: [11]). We extracted DNA from each sample, generated amplicon libraries targeting the bacterial 16s rRNA V4 region, and multiplexed these libraries for sequencing using a common 16s profiling method [12]. Following quality filtering, OTU abundances were estimated using open-reference OTU picking in QIIME [13]; Supplementary Information). We eliminated one ethanol sample because it generated very few reads; all remaining samples were rarefied to 54,633 reads for subsequent analyses. We identified 21,006 OTUs overall (mean per sample=1,656 ±237 s.d.; Table S1).

The resulting data recapitulated previous observations showing storage condition effects on mean alpha diversity (e.g., [2-4, 6]. In our case, samples stored in ethanol, then lyophilized, exhibited lower Shannon’s Diversity Index (SDI) values relative to other conditions (Tukey’s HSD: p between 7.6x10^-5^ and 0.063), and samples stored in RNAlater exhibited somewhat higher values, although this comparison was only significant in comparison to the lyophilized condition (p=7.6x10^-5^; **Figure 1a;** Table S2). However, in spite of these differences, SDI values retained a strong signature of individual identity. Specifically, SDI values were significantly correlated across samples from the same individual across storage methods **(Figure 1b).** Further, although we observed several rank changes across conditions **(Figure 1a)**, individual identity explained a larger proportion of variance in SDI across samples than storage conditions (ANOVA: 50% versus 36%). We obtained qualitatively similar but weaker results for the number of OTUs identified in each sample (Figure S1), suggesting that the combination of species richness and evenness captured by SDI is more stable than richness alone.

**Figure 1.**
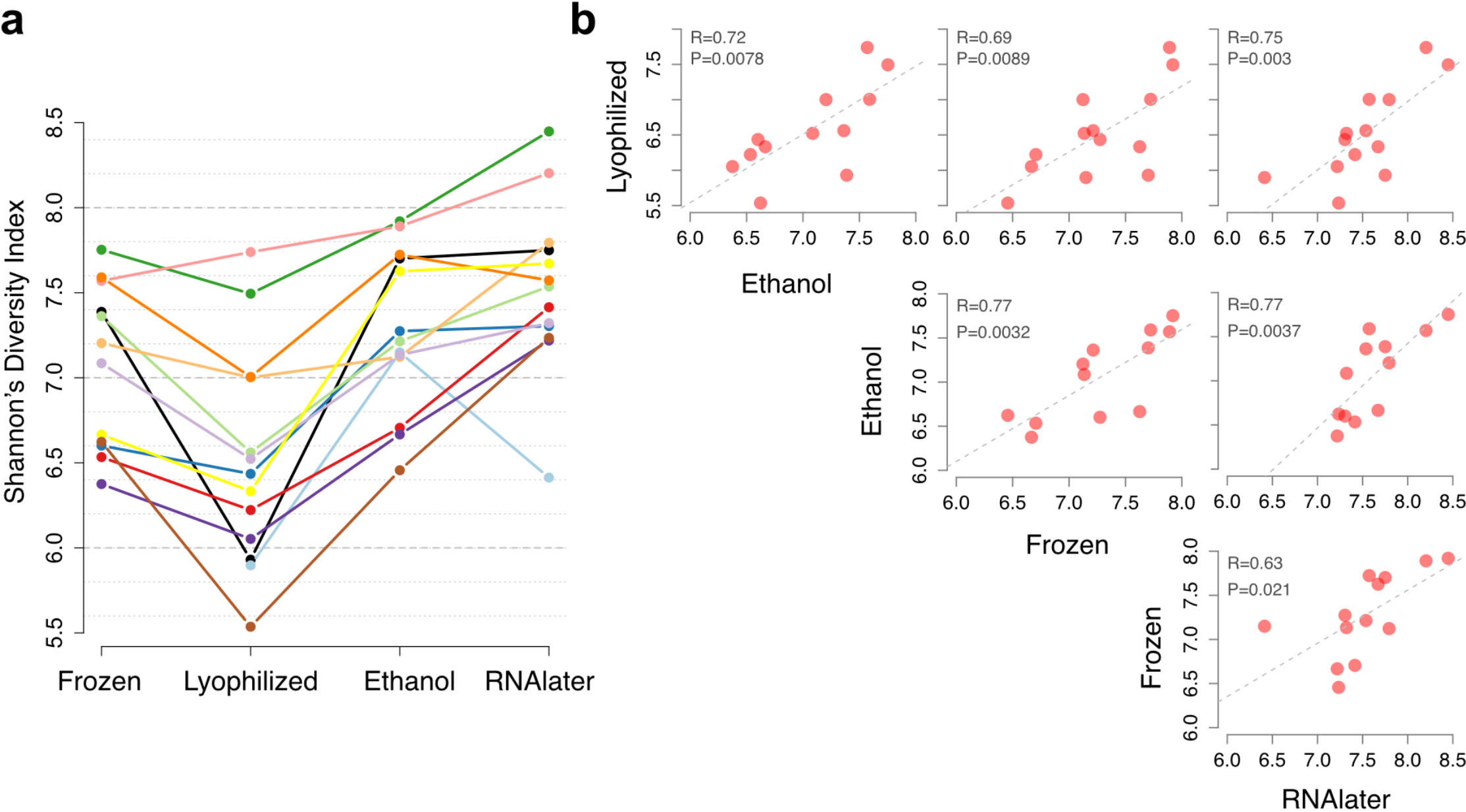
Shannon’s Diversity Index (SDI) values (y-axis) shown as a function of storage method (x-axis), with each individual plotted in a different color. Samples collected in ethanol, then lyophilized, have significantly lower SDI values than other storage methods (Tukey’s HSD p-values range from 7.6 x 10^-5^ to 0.063 across comparisons against lyophilized samples; Table S2). **(b)** SDI values are significantly correlated within individuals, between all storage methods (Pearson’s correlation). Each dot represents an individual, and each panel shows the correlation between SDI values obtained from two different storage methods.

**Figure S1.**
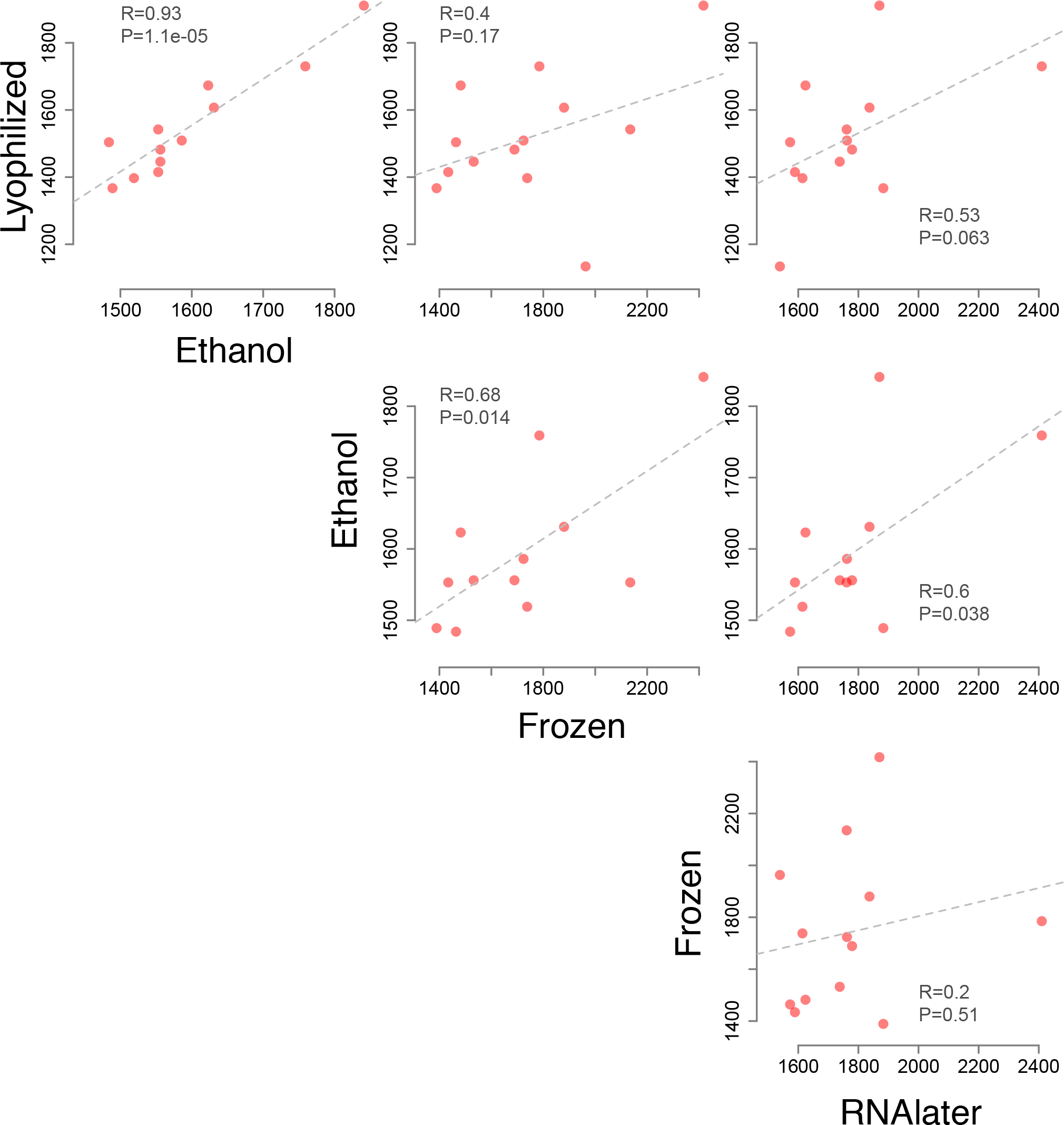
Pairwise correlations for species richness across storage conditions. Correlation vithin individual, between storage methods, for the number of OTUs detected in each sample Pearson’s correlation). Each dot represents an individual, and each panel shows the correlation •etween values in two different storage methods.

Beta diversity measures of community similarity (Tables S3-S5) were also more influenced by individual identity than sample storage condition. 66.3% of variation in taxonomic abundance could be explained by individual identity, compared to 14.3% by storage method (PERMANOVA on a Bray-Curtis dissimilarity matrix; p<0.001 for both predictors). Bray-Curtis dissimilarities were much higher for pairs of samples collected from different individuals (mean=0.51 ±0.11 s.d. within condition; 0.56 ±0.11 s.d. between conditions) than for samples collected from the same individual using different storage conditions (mean=0.35 ±0.11 s.d., **Figure 2a;** see also Figure S2). Samples from the same individual, but not storage condition, also clustered together in a hierarchical clustering analysis using either Bray-Curtis or unweighted UniFrac measures **(Figure 2b;** Figure S3a; no clustering, either by individual or storage condition, was observable using weighted UniFrac: Figure S3b). Most importantly, relative distances between individuals remained consistent across storage conditions. For example, pairwise correlations between Bray-Curtis dissimilarity matrices calculated separately for each condition were highly correlated (r=0.59 to 0.88, all p<0.005; Table S6), with similar patterns observed using weighted or unweighted UniFrac (Tables S7-S8).

**Figure 2.**
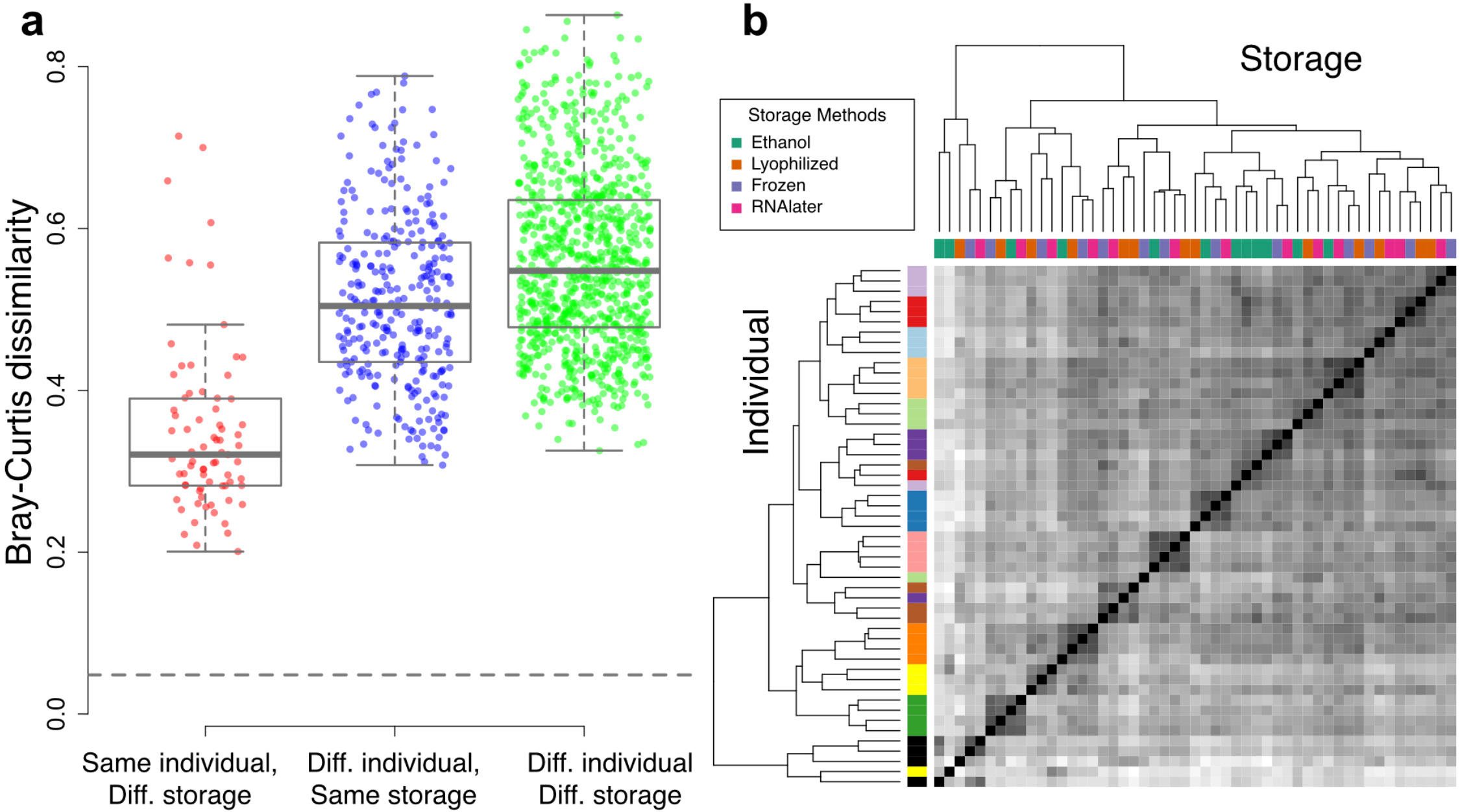
Bray-Curtis dissimilarity values (y-axis) comparing the same individual from samples collected under different storage conditions (red), different individuals with samples collected under the same storage conditions (blue), and different individuals with samples collected under different storage conditions (green). Median Bray-Curtis dissimilarity calculated from subsampling reads from the same sample (i.e., the minimum dissimilarity due to read resampling alone) is indicated by the gray dashed line. Because of the large number of data points, all pairwise comparisons are highly significant (Wilcoxon Rank Sum test, p<1 x 10^-9^). However, the dissimilarity values for same individual/different storage are much lower on average (mean=0.35 ±0.11 s.d.) than dissimilarity values measured between individuals in either the same (mean=0.51 ±0.11 s.d.) or different (0.56 ±0.11 s.d.) storage conditions. **(b)** Bray-Curtis dissimilarities cluster more strongly by individual (colors along the left-hand sidebar, with one color per individual) than by storage method (colors shown on the top, next to the dendrogram, and in the boxed legend).

**Figure S2.**
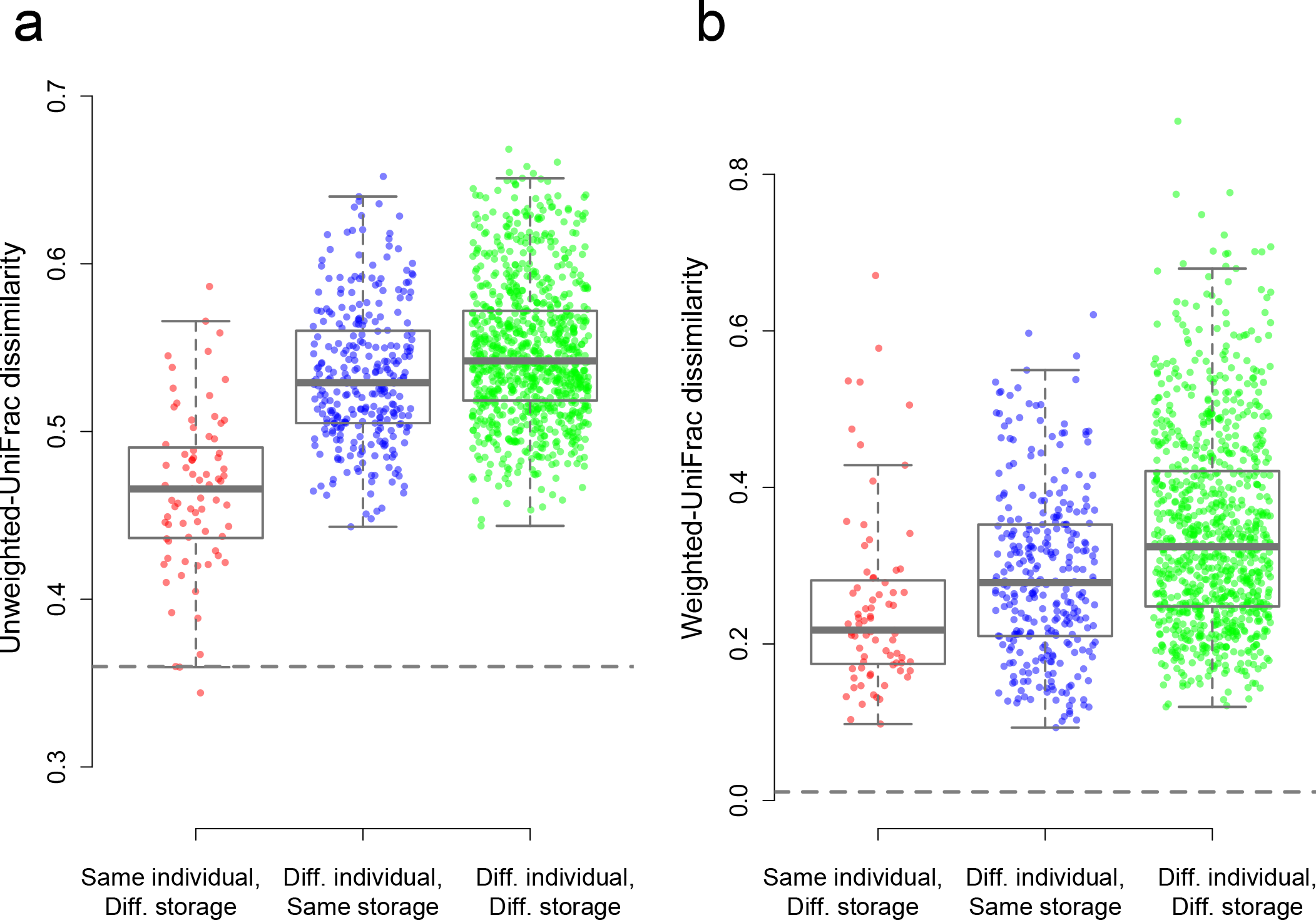
Weighted and unweighted UniFrac dissimilarity values between different sample types. (A) Weighted UniFrac dissimilarities comparing the same individual from amples collected under different storage conditions (red), different individuals with samples collected under the same storage conditions (blue), and different individuals with samples collected under different storage conditions (green). (B) As in A, but for unweighted UniFrac lissimilarities. In both A and B, the grey dashed line indicates the median dissimilarity value calculated from subsampling reads from the same sample (i.e., the minimum dissimilarity due to ead resampling alone).

**Figure S3.**
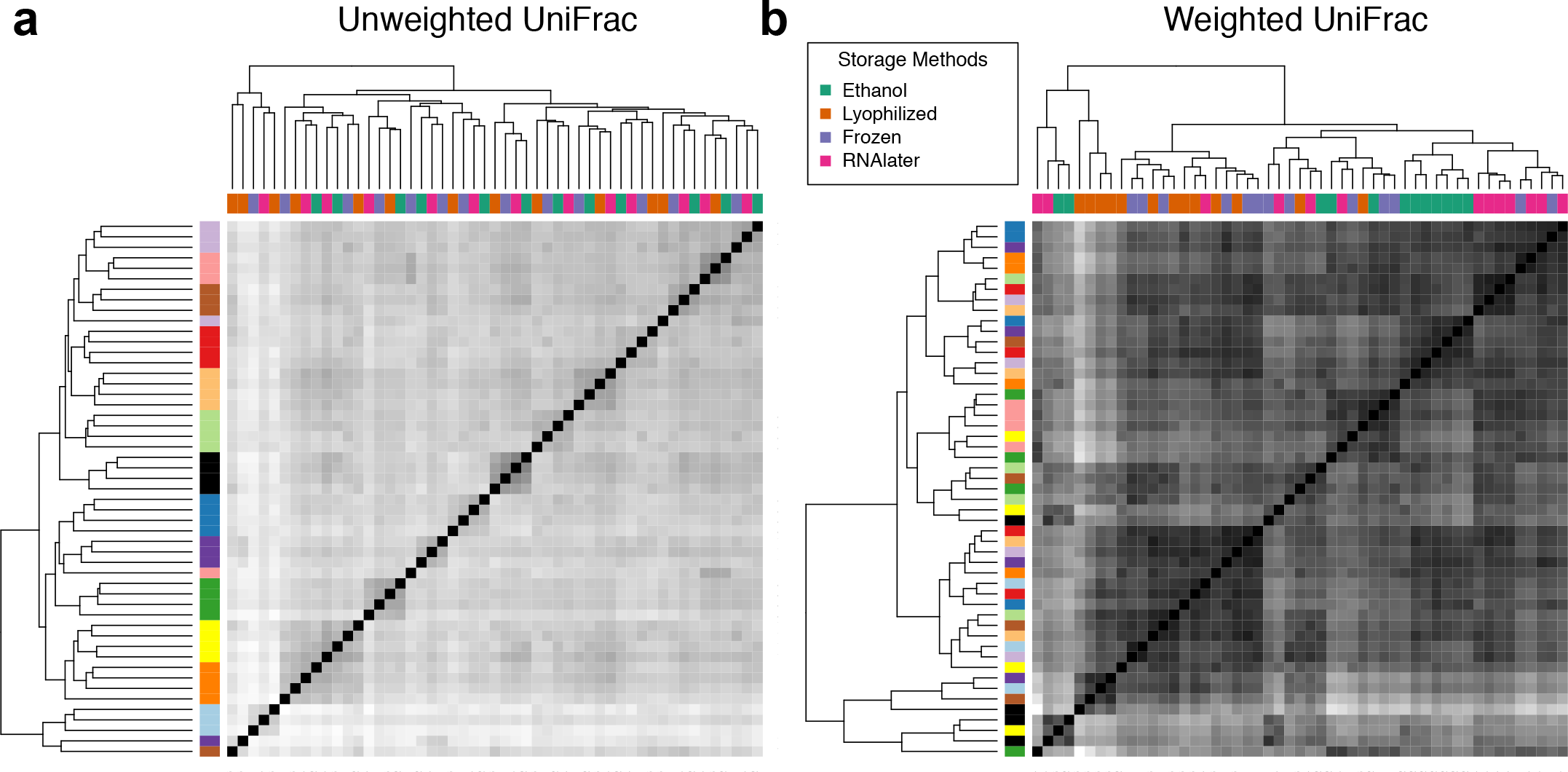
Hierarchical clustering plots based on unweighted and weighted UniFrac dissimilarity measures. Like Bray-Curtis dissimilarities (main text, Figure 2), (A) unweighted JniFrac dissimilarities cluster more strongly by individual (colors along the lefthand sidebar, vith one color per individual) than by storage method (colors shown on the top). (B) Weighted niFrac similarities show no clear clustering by individual or storage method.

Together, our results indicate that, while mean alpha and beta diversity values are sometimes altered by storage condition, biologically relevant signatures of individual identity tend to be retained, especially for measures of beta diversity. Our findings agree with previous studies using fewer individuals [5-7, 9], and extend them to three of the most commonly used storage methods in vertebrate field studies. For many types of studies, storage condition *per se* may therefore be less important than maintaining consistency in storage methods within a data set. Indeed, our estimates of the effects of individual identity are probably conservative given the standardized housing, diet, and social group structure of our study subjects.

Our findings thus indicate the utility of using banked fecal sample collections from field studies for analyses of gut microbiome variation. These collections are not only substantial (ranging up to tens of thousands of samples), but are also often longitudinal, complemented by extensive demographic and behavioral metadata, and focused on species of particular conservation concern. As such, they represent extraordinary, largely untapped resources for understanding the causes, consequences, and diversity of gut microbial structure.

## Acknowledgements

We gratefully acknowledge support from the National Science Foundation (IOS 1053461 to E.A.A.), National Institutes of Health (R01-GM102562 to J.T., L.B.B., and M.E.W.), and the Clare Boothe Luce Foundation (E.A.A.). We also thank A. Tripp, J. Whitley, and J. Johnson for their support in collecting samples, M. Gearhart for assistance with computational analysis, and members of the Blekhman lab for helpful discussions. This work was carried out using computing resources at the Minnesota Supercomputing Institute.

## Availability of data and materials

All sequencing data will be made available in the NCBI Short Read Archive upon publication.

## Supplementary Information for Blekhman et al, Common methods for fecal sample storage n field studies yield consistent signatures of individual identity in microbiome sequencing data

**1. Supplementary Methods**

A. Study subjects and sample collection
B. 16s rRNA sequencing
C. Low level data processing and OTU table construction
D. Alpha and beta diversity analyses
E. Rarefaction analysis

**2. Supplementary Tables**

Supplementary Tables are provided as a single.xls file, with the following spreadsheets:

**Table S1.** Sample information and summary of sequencing results
**Table S2.** Differences in mean SDI across storage conditions
**Table S3.** Bray-Curtis dissimilarity values between all samples
**Table S4.** Unweighted UniFrac distances between all samples
**Table S5.** Weighted UniFrac distances between all samples
**Table S6.** Correlations between Bray-Curtis dissimilarity matrices obtained from different storage conditions
**Table S7.** Correlations between Weighted UniFrac dissimilarity matrices obtained from different storage conditions
**Table S8.** Correlations between Unweighted UniFrac dissimilarity matrices obtained from different storage conditions.

**3. Supplementary Figures**

**Figure S1.** Pairwise correlations for species richness across storage conditions
**Figure S2.** Weighted and unweighted UniFrac dissimilarity values between different sample types
**Figure S3.** Hierarchical clustering plots based on unweighted and weighted Unifrac dissimilarity measures
**Figure S4.** Statistical robustness to rarefaction

## A. Study subjects and sample collection

Study subjects were 13 adult female rhesus macaques (*Macaca mulatta*), members of 7 lifferent social groups housed at the Yerkes National Primate Research Center (YNPRC). These groups were formed as part of a separate study on the relationship between dominance rank and cene regulation. All groups were maintained in standardized indoor-outdoor housing runs (25 m : 25 m per run), under standardized demographic (5 adult females per group), dietary, and observational conditions.

Fecal samples were collected opportunistically, within 10-15 minutes after deposition, vhen females were briefly separated from the rest of their social groups for other purposes. Each sample was subdivided into four equal subsamples, with the first subsample frozen immediately at −20 °C; the second subsample immersed in the commercial preservative RNAlater (Life Technologies, Carlsbad, CA); and the third and fourth subsamples immersed in absolute ethanol. Samples were shipped overnight to Duke University either on dry ice (immediately frozen samples) or at room temperature (RNAlater and ethanol samples). At Duke, one of the ethanol-tored subsamples was processed following standard methods used to process fecal samples for teroid hormone analysis in primate field studies [1]; see also https://amboselibaboons.nd.edu/assets/75656/altmannlabprotocolsjan08.pdf). In brief, the ethanol storage medium was evaporated under a fume hood. The resulting dried sample was then lyophilized at −50 °C under 0.1 millibar of vacuum pressure, and then sifted through a fine mesh strainer to separate fecal matter from large, undigested fragments of vegetation.

DNA from all samples (n=52, representing 4 storage conditions for 13 individuals) was ¡xtracted using MO BlO’s PowerSoil DNA Isolation kit (MO BIO Laboratories, Inc., Carlsbad, ^A). Extractions were conducted according to the manufacturer’s instructions, except that for lyophilized samples, extractions were obtained from 0.05 g of sample instead of 0.25 g to avoid complete absorption of liquid in the first steps of the DNA extraction, and subsequent extraction ailure.

## B. Ì. 16s rRNA sequencing

Purified DNA samples were shipped to the University of Minnesota Genomics Center for library preparation and sequencing. DNA isolated from fecal samples was quantified by 16S rRNA sequencing. The V4 region of the 16S rRNA gene was PCR amplified, using forward and everse primers 515F

TCGTCGGCAGCGTCAGATGTGTATAAGAGACAGGTGCCAGCMGCCGCGGTAA) and 06R

GTCTCGTGGGCTCGGAGATGTGTATAAGAGACAGGGACTACHVGGGTWTCTAAT) 2], followed by amplification with indexing primers (forward: ^ATGATACGGCGACCACCGAGATCTACAC[i5]TCGTCGGCAGCGTC, reverse::AAGCAGAAGACGGCATACGAGAT[i7]GTCTCGTGGGCTCGG, where [i5] and [i7] refer to the index sequence codes used by Illumina). PCR amplification with the 515F/806R primer >air was conducted using KAPA HiFidelity Hot Start Polymerase, under the following conditions: an initial denaturation step at 95 °C for 5 min, followed by 20 cycles of denaturation 20 s at 98 °C), annealing (15 s at 55 °C), and elongation (60 s at 72 °C). Amplified samples vere diluted 1:100 in water, and 5 μL of the 1:100 diluted sample were used for the second PCR reaction with the indexing primers, using the same cycling conditions but for 10 cycles instead of ‘0. Pooled and size-selected samples were denatured in NaOH, diluted to 8 pM in Illumina’s ÎT1 buffer, spiked with 15% PhiX, and heat denatured at 96 °C for 2 minutes immediately prior to loading on a MiSeq flowcell. We produced 300 bp paired-end sequences, with a mean of ^38,734 (±131,840 s.d.) fragments sequenced for each sample (see Table S1 for sample-specific information).

## C. Low level data processing and OTU table construction

Following sample demultiplexing, primer sequences were removed from the raw reads ising CutAdapt v.1.7.1 [3]. Because CutAdapt does not always detect reverse primers effectively, the first 29 base pairs (theoretically the primer and linker sequences) were removed from reverse reads. Reads were truncated at the first base pair with a PHRED quality score ≤3, and forward md reverse reads were then merged using USEARCH v6.1 [4]. Read pairs that failed to merge vere discarded. We used QIIME v1.8 to conduct further quality control filtering [5]. QIIME was run with default parameters except for the minimum acceptable per-base Phred score parameter, which we increased from 4 to 20. Putative chimeric sequences were identified using UCHIME implemented in USEARCH v6.1) [6], and sequences were discarded from the sample when both reference-based (against the RDP Gold training database v9: [7] and *de novo* abundance-based nethods flagged them as likely chimeras. Chimeric sequences constituted 0.009% of our sequencing reads.

To identify operational taxonomic units (OTUs) in our data set, we used the open-eference OTU picking pipeline in QIIME. Specifically, the set of chimera-filtered reads was first clustered using the UCLUST v1.2.22 algorithm and the GreenGenes database (May 2013 release: [8], with a 97% identity threshold. Sequences that failed to cluster against the reference database were then clustered *de novo*, with sequences that failed both clustering attempts discarded. A representative sequence for each cluster was selected based on the most abundant sequence, and then aligned using PyNAST v1.2.2 [9]. Sequences that failed to align were liscarded. Taxonomic identity was assigned to aligned OTUs using the RDP classifier v2.2 [10], retrained to the May 2013 release of the GreenGenes database [8]. Singleton OTUs were emoved from the OTU table as they tend to be enriched for sequencing errors. At this stage, we also removed one sample, the ethanol-stored sample for individual Ia13, due to low read count 629 reads following quality control filtering).

For all subsequent analyses, we rarefied the OTU table to 54,633 reads per sample using he QIIME v 1.8.0 script singlerarefaction.py. Subsampling reads from individuals with iniformly high coverage across storage conditions supported the stability of our summary statistics at this level of rarefaction (see Supplementary Methods, section E).

## D. Alpha and beta diversity analyses

We calculated alpha diversity measures (Shannon’s Diversity Index and number of »bserved OTUs as a measure of species richness) using the QIIME v 1.8.0 script alpha_diversity.py, and all beta diversity measures (Bray-Curtis dissimilarity, unweighted JniFrac, and weighted UniFrac) using the corresponding QIIME v 1.8.0 script betadiversity.py 5]. To estimate the minimum beta diversity dissimilarity due to random resampling error, we ised high sequencing depth samples for which at least five times the number of rarefied reads vere available (273,165 reads; n = 12). We drew five random subsamples from the total quality-iltered read count for each of these samples. We then calculated all pairwise Bray-Curtis lissimilarity values between subsamples from the same original sample. The median of these dissimilarity values is shown as the dashed line in Figure 2a (and for weighted and unweighted UniFrac analyses in Figure S2).

All statistical analyses on alpha and beta diversity values were conducted in R v 3.1.1 [11] ising either the R base packages or, for PERMANOVA, the R package *vegan* [12] and the R lackage *ade4* v 1.7-2 [13].

## E. Rarefaction analysis

After rarefaction, we retained a read depth of 54,633 reads per sample. To test whether this read depth affected our ability to estimate correlations between SDI or Bray-Curtis dissimilarities across storage conditions, we investigated the relationship between read depth, SDI, and the dissimilarity values using the five individuals for whom we produced a large lumber of reads across all four storage conditions (n=20 samples for Ve12, Tf12, Pp10, Js11, and Jj10). For these individuals, we computed the SDI and Mantel test correlation between Bray-turn’s dissimilarities at 10 different rarefaction levels: 98,039 reads, 54,633 reads (corresponding to the sequencing depth used in the main analyses), 27,326 reads, 13,658 reads, 6,829 reads, 3,414 reads, 1,707 reads, 854 reads, 427 reads, and 213 reads. We performed a similar analysis for the Wilcoxon rank sum statistic comparing Bray-Curtis dissimilarities from samples taken from the same individual, stored in different conditions, to samples taken from different individuals, from the same storage condition. In all cases, estimates stabilized at sequencing lepths lower than the one used in the main analysis (i.e., 54,633 reads), suggesting that our malyses are not influenced by coverage concerns (Figure S4).

**Figure S4.**
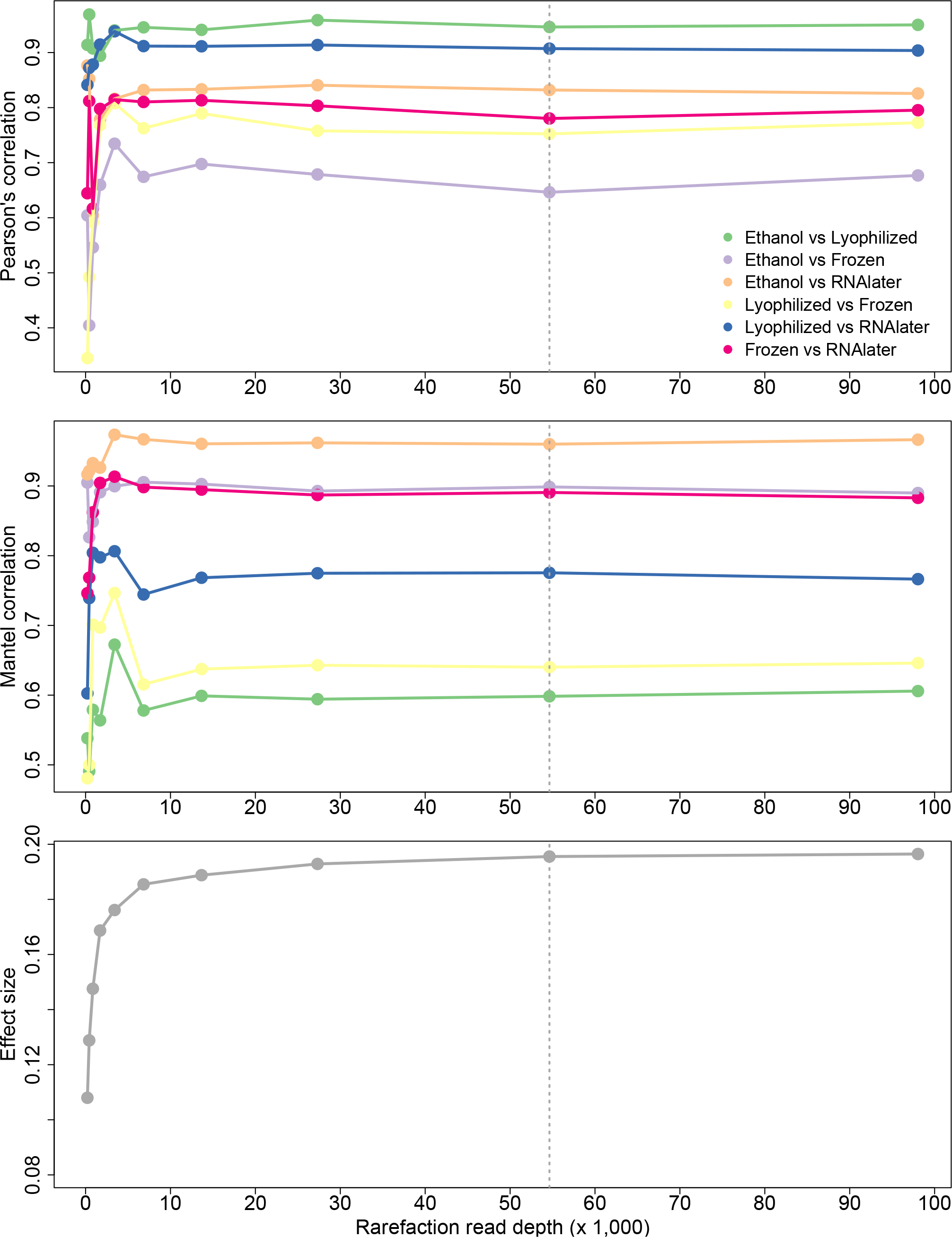
Statistical robustness to rarefaction. We used samples from 5 individuals with high coverage across all four storage conditions to test whether the rarefied sequencing coverage used n our main analysis (gray dashed line) was sufficient to produce stable estimates of (i) SDI correlations across storage condition (top panel); (ii) Mantel test correlations of Bray-Curtis issimilarity values across storage conditions (middle panel); and (iii) Wilcoxon test statistics for the difference in mean pairwise Bray-Curtis dissimilarity values when comparing samples from he same individual stored in different conditions to samples from different individuals stored in the same condition (bottom panel). In all three cases, test statistics stabilize by a depth of 20,00010,000 rarefied reads, lower than the read depth used in our actual analyses.

